# DIA-based systems biology approach unveils novel E3-dependent responses to a metabolic shift

**DOI:** 10.1101/2020.09.28.315465

**Authors:** Ozge Karayel, André C. Michaelis, Matthias Mann, Brenda A. Schulman, Christine R. Langlois

## Abstract

Yeast *Saccharomyces cerevisiae* is a powerful model system for systems-wide biology screens and large-scale proteomics methods. Nearly complete proteomics coverage has been achieved owing to advances in mass spectrometry. However, it remains challenging to scale this technology for rapid and high-throughput analysis of the yeast proteome to investigate biological pathways on a global scale. Here we describe a systems biology workflow employing plate-based sample preparation and rapid, single-run data independent mass spectrometry analysis (DIA). Our approach is straightforward, easy to implement and enables quantitative profiling and comparisons of hundreds of nearly complete yeast proteomes in only a few days. We evaluate its capability by characterizing changes in the yeast proteome in response to environmental perturbations, identifying distinct responses to each of them, and providing a comprehensive resource of these responses. Apart from rapidly recapitulating previously observed responses, we characterized carbon source dependent regulation of the GID E3 ligase, an important regulator of cellular metabolism during the switch between gluconeogenic and glycolytic growth conditions. This unveiled new regulatory targets of the GID ligase during a metabolic switch. Our comprehensive yeast system read-out pinpointed effects of a single deletion or point mutation in the GID complex on the global proteome, allowing the identification and validation novel targets of the GID E3 ligase. Moreover, our approach allowed the identification of targets from multiple cellular pathways that display distinct patterns of regulation. Although developed in yeast, rapid whole proteome-based readouts can serve as comprehensive systems-level assay in all cellular systems.

## INTRODUCTION

Proteome remodeling has repeatedly proven to be a vital cellular mechanism in response to stress, changes in environmental conditions, and toxins or pathogens. Cells must both synthesize proteins which enable them to adapt to the new environmental condition, and inactivate or degrade proteins which are detrimental or no longer needed. For each environmental perturbation, the proteome must be precisely and distinctly remodeled to ensure healthy and viable cells (1). Indeed, decreases in proteome integrity are hallmarks of many human diseases, including cancer, Alzheimer’s disease, muscular dystrophies, and cystic fibrosis (2-4). Despite the importance of cellular stress response, our understanding of how cellular pathways interact during adaptation remains incomplete. Therefore, knowing precisely how the proteome changes at a global level in response to environmental cues is crucial for identifying the underlying molecular mechanisms that facilitate cellular adaptation.

Yeast *Saccharomyces cerevisiae* is a powerful model system that is widely used to probe biological pathways, due to its ease of manipulation and rapid growth compared to mammalian models. In addition, the availability of extensive genetic resources in yeast, including deletion libraries (5, 6), GFP-tagged libraries (7, 8), and over-expression libraries (9) and the recently developed SWAp-tag library (10-12) has made yeast a premier model system for conducting transcriptomics, proteomics, interactomics or metabolomics screens (13-19). Indeed, systems-wide biology screens and large-scale proteomics were both pioneered in the yeast model. Furthermore, the cellular interaction networks and molecular mechanisms ascertained in yeast can be readily applied to other systems (20-22).

Early genome-wide studies showed that over 4,000 proteins are expressed during log-phase growth in yeast and this organism was the first whose entire proteome was mapped by mass spectrometry (MS)-based proteomics (23). Subsequently, yeast has served as a model of choice for the development of ever more sensitive and faster proteomics workflows (23-34). Remarkably, the optimized sample preparation coupled with MS analysis performed on the Orbitrap hybrid mass spectrometer allowed identification of around 4,000 yeast proteins over a 70-min LC-MS/MS run (24, 30). However, the necessity of technological expertise and lengthy analysis times for high-quality, in-depth yeast proteome measurements has so far precluded the widespread adoption of cutting-edge proteomics workflows in the yeast research community. With further advances in technology and new acquisition modes, such as data-independent acquisition (DIA) (35, 36) we hypothesized that it would now be possible to obtain accurate and high yeast proteome coverage by a straightforward and rapid single run approach, enabling researchers to easily study biological processes on a global scale. Such a system could then serve as a template for more complex proteomes, including the human proteome.

One mechanism of maintaining proteome integrity is the marking and degradation of proteins that are damaged or no longer needed with ubiquitin. The conjugation of ubiquitin to its targets is catalyzed by E3 ubiquitin ligases, a diverse group of enzymes that recognize and bind target proteins, and facilitate ubiquitin transfer together with an E2, ubiquitin conjugating enzyme. Ubiquitination relies on a variety of cellular signals to direct E3 ligases to their target proteins, and tight regulation of this process is crucial for cellular viability (37). For instance, during carbon starvation, yeast cells induce expression of the inactive GID (*<u>g</u>*lucose-*<u>i</u>*nduced *<u>d</u>*egradation) E3 ligase, which is subsequently activated upon glucose replenishment. Following its activation, the GID E3 ligase targets gluconeogenic enzymes, leading to their degradation, and allowing the yeast cell to proceed with aerobic growth (38-40). In addition, ubiquitin ligases also serve as crucial regulators in response to oxidative, heavy metal, and protein folding stresses (41-43). Despite the importance of ubiquitination during cellular adaptation, our knowledge of the E3-dependent responses to cellular perturbation remains incomplete.

Here, we describe a systems biology approach employing rapid, single-run data independent (DIA) mass spectrometric analysis, which we use to comprehensively map changes to the yeast proteome in response to a variety of yeast stresses. We investigate growth conditions commonly used in yeast research, including growth media, heat shock, osmotic shock, amino acid starvation, and nitrogen starvation. Our DIA-based approach is sufficiently sensitive and robust to detect quantitative proteome remodeling in response to all these stresses. We then apply our methodology to probe a specific biological question to identify novel regulation by the GID E3 ligase during a metabolic switch. We use a combination of a core subunit deletion and a structure-based catalytic mutant to identify all the known substrates of the GID E3 ligase, and discover two previously unknown targets which display distinct patterns of regulation.

## RESULTS

### Streamlined and scalable yeast proteome analysis employing data-independent acquisition

In order to establish a fast and scalable single run analysis approach for yeast proteome profiling, we explored a data-independent acquisition (DIA) strategy on an Orbitrap mass spectrometer. Unlike data-dependent acquisition (DDA), a DIA method isolates co-eluting peptide ions together in predefined mass windows, fragmenting and analyzing all ions simultaneously (36). This strategy overcomes the limited sequencing speed of sequential DDA, enabling fast and scalable single-shot analysis workflows. On Orbitrap-based mass analyzers it yields substantially higher number of identified proteins with unprecedented quantitative accuracy (44). To generate a yeast-specific and comprehensive spectral library that is generally used for this approach, we cultured yeast under various growth and stress conditions. After extraction and digestion of proteins, we separated peptides obtained from each condition by basic reversed-phase (bRP) chromatography into 8 fractions (**Figure 1A**). The resulting 64 fractions (8 fractions x 8 conditions) were measured using a DDA method with 23 minutes LC gradient and analyzed with the Spectronaut software. Together with LC overhead time this took about half an hour, allowing for the analysis of 45 samples per day – almost half a 96 well plate. Our library comprised more than 74,103 precursors which mapped into 4,712 unique proteins, covering 87% of the expressed yeast proteome according to a previous report that computationally aggregated 21 different large-scale datasets (45).

**Figure 1.**
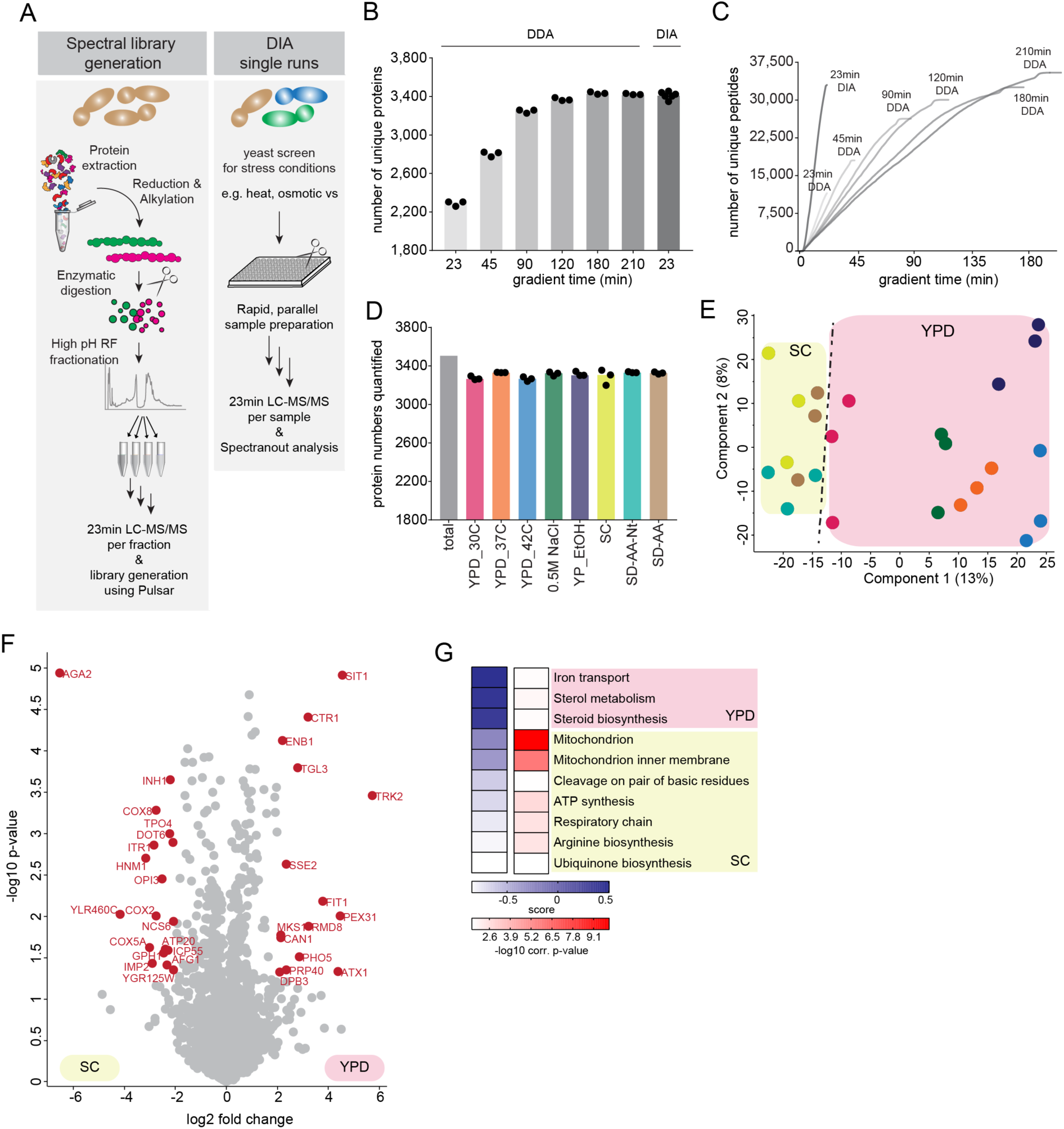
Fast and scalable yeast proteome analysis using data-independent acquisition. **A**. Experimental workflow for yeast spectral library construction (left panel) and fast, single run DIA-based analysis of yeast proteomes (right panel). **B**. Number of identified proteins using DDA with varying LC gradient lengths compared to 23 minutes DIA. **C**. Cumulative number of identified unique yeast peptides over time using DDA with varying LC gradient lengths and the 23 minutes DIA method. **D**. Number of quantified proteins in growth and stress conditions. **E**. PCA of conditions along with their biological replicates based on their proteomic expression profiles. **F**. Volcano plot of the (-log10) p-values vs. the log2 protein abundance differences between yeast grown in YPD vs. SC. The proteins marked in red change significantly (p value < 0.05 and at least 4-fold change in both directions). **G**. GO-term enrichment in the YPD vs. SC fold change dimension (1D enrichment, FDR < 5%). Terms with positive enrichment scores are enriched in YPD over SC and vice versa.

Combined with our own comprehensive spectral library, the 23 minutes DIA method on average identified 33,909 peptides and 3,413 distinct proteins in single measurements of six replicates (Q-value less than 1% at protein and precursor levels, **Figure 1B-C, Supplementary Table 1**). This implies that approximately 73% of proteins in the deep yeast spectral library were matched into the single runs. Note that the library combines many different proteome states and the single runs represent only one, so the degree of proteome completeness is likely much higher than 73%. Measurements were highly reproducible with Pearson coefficients greater than 0.92 between replicates (**Figure S1A**) and CVs < 20% for 68% of all common proteins between the six replicates. In comparison, a single run data-dependent acquisition strategy with the same LC gradient quantified only 11,883 peptides and 2,289 distinct proteins on average (**Figure 1B-C**). To more directly compare the performance of the 23-minute DIA method to the DDA method we analyzed the same sample with increasing gradient lengths. We could only reach the same depth using DDA method with at least 180-minute-long LC gradients (33,425 peptides and 3,435 proteins) (**Figure 1B-C**). Thus, the DIA method allows us to obtain comparable coverage to DDA in high throughput and in-depth fashion in almost one-tenth of the analysis time.

**Table 1:**
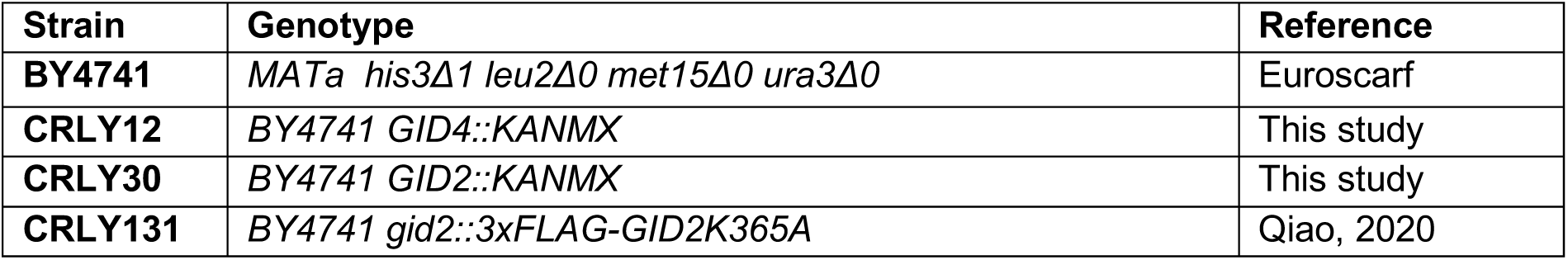
Yeast strains used in this study

| Strain | Genotype | Reference |
| --- | --- | --- |
| **BY4741** | *MATa his3Δ1 leu2Δ0 met15Δ0 ura3Δ0* | Euroscarf |
| **CRLY12** | *BY4741 GID4::KANMX* | This study |
| **CRLY30** | *BY4741 GID2::KANMX* | This study |
| **CRLY131** | *BY4741 gid2::3xFLAG-GID2K365A* | Qiao, 2020 |

### Large-scale and quantitative analysis of yeast stress response in half a day

Using our DIA-based systems biology approach, we next comprehensively and quantitatively analyzed proteome changes in response to various stresses in yeast. Each condition was processed in three biological replicates and – after tryptic digestion – the peptides were analyzed in single runs using our rapid DIA method. We quantified 3,506 distinct proteins in total (**Figure 1D, Supplementary Table 2**). Reproducibility was high, with Pearson correlations > 0.93 between the three biological replicates (**Figure S1B**). Strikingly, over 90% of all detected proteins were consistently quantified at varying levels across all conditions (**Supplementary Table 2**). Principal component analysis (PCA) demonstrated that the first component accounted for 13% of the variability and segregated with the different conditions and growth media as the major effectors (**Figure 1E**).

**Table 2:** Plasmids used in this study

| Plasmid |  | Reference |
| --- | --- | --- |
| **CRLP47** | pRS313-P<sub>TDH3</sub>(modified)-Aro10<sub>3xFlag</sub>-CYC-p<sub>TDH3</sub>(modified)-<br>flagDHFR<sub>ha</sub>-CYC | This study |
| **CRLP48** | pRS313-P<sub>TDH3</sub>(modified)-Acs1<sub>3xFlag</sub>-CYC-p<sub>TDH3</sub>(modified)-<br>flagDHFR<sub>ha</sub>-CYC | This study |

We first looked more closely at the differences in protein expression during growth in YPD (yeast extract peptone dextrose; rich media) and SC (synthetic complete media), the two most common growth media used in yeast research. YPD and SC media differ in their nutrient composition, as well as their pH. During growth in YPD, the three most significantly upregulated proteins (Sit1, Ctr1, and Enb1) are regulators of copper and iron transport (**Figure 1F (right),1G**), consistent with the fact that copper and iron are limiting factors for the growth of yeast at more alkaline pH (46). Conversely, during growth in SC, many mitochondrial proteins were upregulated compared to YPD (**Figure 1F (left), 1G**), including the cytochrome c oxidase subunits Cox8, Cox2 and Cox5a, the mitochondrial ATP synthase, Atp20, and the mitochondrial aminopeptidase, Icp55. Yeast mitochondria reproduce through fission and must be inherited by daughter cells during cell division (47). The upregulation of many mitochondrial proteins is thus consistent with the faster growth rate of our yeast strains in SC compared to YPD. Because the choice of media is often considered crucial in experimental design, this data on differentially regulated proteins in pathways of interest provide an important resource for yeast biologists.

Next, we investigated proteome changes in yeast grown under various stress conditions. Here, we focused on those commonly utilized in yeast research: heat shock, osmotic shock, carbon starvation, amino acid starvation, and nitrogen starvation. Each produced a discrete stress response, resulting in synthesis or degradation of a distinct set of proteins (**Figure 2A, S2A-F**). For example, yeast cells grown under heat shock induce expression of chaperones and stress response proteins, a well-characterized response that allows the cell to quickly recover from global heat-induced protein misfolding (48-50). Importantly, our data also revealed that the heat shock response is dose dependent, with higher induction of the stress response at 42°C compared to 37°C (**Figure 2A (teal cluster), 2B**). Yeast experiencing osmotic shock, on the other hand, induced distinct proteome changes, with the most enriched GO-term under this condition being actin-cortical patch (**Figure S2A-B**). This is consistent with the fact that yeast cells rapidly disassemble and remodel the actin cytoskeleton during osmotic stress and favor the formation of actin patches over filaments, a mechanism that lowers the turgor pressure and allows continued growth of yeast under high osmolarity (51, 52). In addition, one of the most upregulated proteins during osmotic stress is Ena1 (**Figure S2A**), a sodium efflux pump that plays a crucial role in allowing salt tolerance (53). Growth during amino acid or nitrogen starvation primarily resulted in the induction of amino acid biosynthetic pathways, with arginine and cysteine synthesis being particularly upregulated (**Figure S2C-F**).

**Figure 2.**
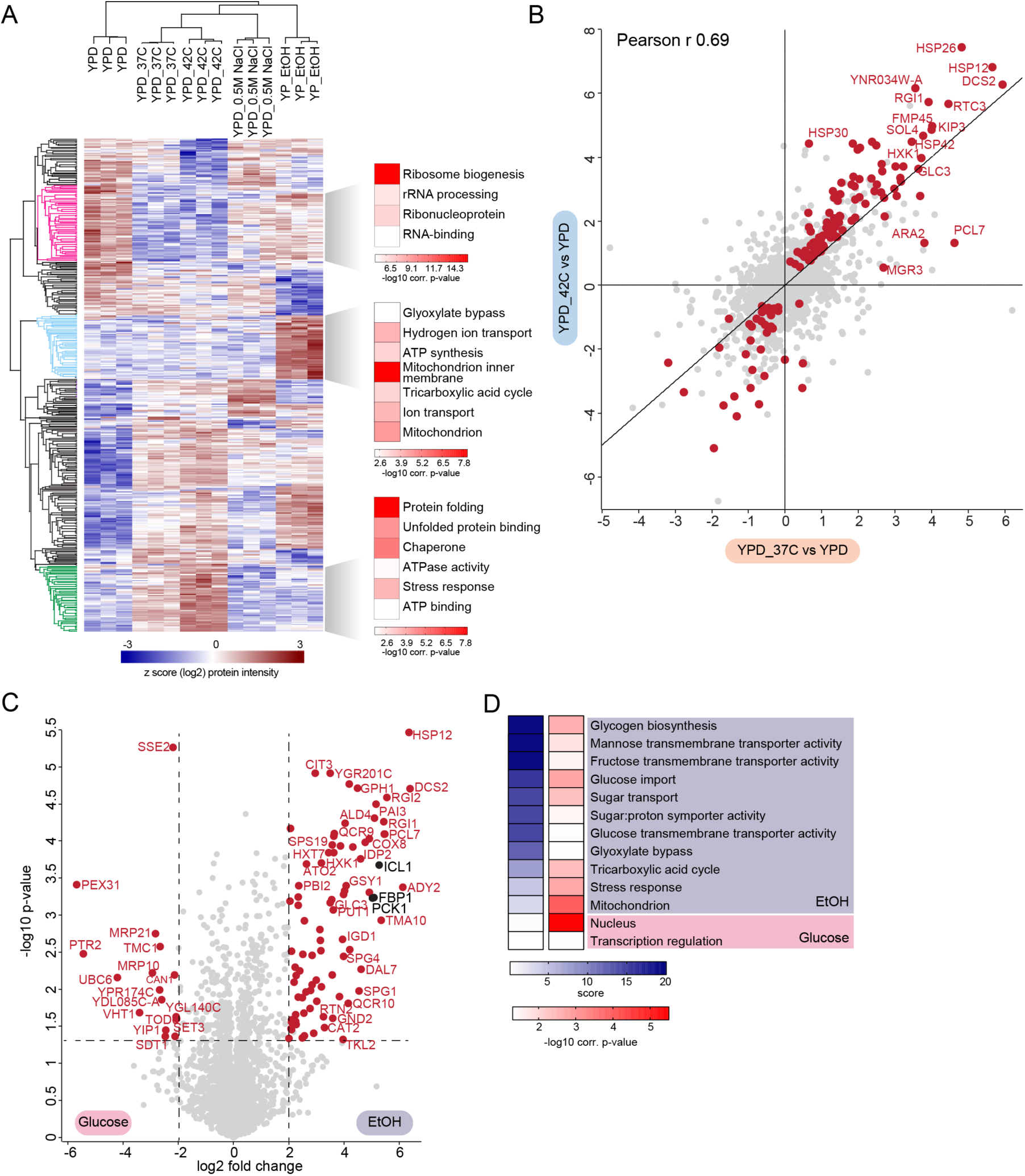
Large-scale and quantitative analysis of yeast proteomes under different stresses. **A**. Heat map of z-scored protein abundances (log2 DIA intensities) of the differentially expressed proteins (ANOVA, FDR<0.01) after hierarchical clustering of stress conditions performed in YPD and YPE. Fisher exact test was performed to identify significantly enriched GO-terms in the most prominent profiles (FDR < 5%). **B**. Correlation of log2 fold-changes of the all quantified proteins in HS experiment. The proteins that change significantly in either 37°C or 42°C compared to 30°C YPD control are colored in red (T-test, FDR < 5%). **C**. Volcano plot of the (-log10) p-values vs. the log2 protein abundance differences between glucose starvation (ethanol) vs. YPD. Red dots indicate significantly different proteins, determined based on p value < 0.05 and at least 4-fold change in both directions. **D**. GO-term enrichment in the ethanol vs. YPD fold change dimension (1D enrichment, FDR < 5%). Terms with positive enrichment scores are enriched in stress condition over glucose (YPD) control and vice versa.

In addition to temperature and nutrient availability, carbon source is a crucial determinant of yeast growth. We compared the proteomes of yeast grown in the aerobic carbon source, glucose (YPD), with the non-fermentable carbon source ethanol. Yeast will preferentially metabolize aerobic carbon sources, such as glucose, when they are present in the media. When only non-fermentable carbon sources like as ethanol are present, yeast cells will instead metabolize them through several pathways, including gluconeogenesis to generate glucose, and conversion of ethanol into pyruvate to allow for ATP generation in the mitochondria via the tricarboxylic acid cycle (54, 55). Consistent with this, we observe a general upregulation of mitochondrial proteins, and those involved in the tricarboxylic acid cycle during growth in ethanol (**Figure 2A, light blue cluster, 2C-D**). In addition, many proteins involved in carbon metabolism are differentially regulated in glucose and ethanol containing media. For example, we see a greater than 16-fold upregulation of the gluconeogenic enzymes Fbp1, Pck1, and Icl1 (**Figure 2C**). In the absence of glucose, both Hxt7, a glucose transporter, and Hxk1, a hexokinase, are upregulated (**Figure 2C**), allowing the cell to quickly import and metabolize any glucose in the environment. These results are consistent with the idea that yeast have “anticipatory” programing, which not only allows them to adapt to current stressor, but also facilitates a rapid response to shifts in environmental conditions (56). Moreover, apart from identifying proteins that have altered levels in response to a shift in environmental conditions, we also accurately determined their fold changes, giving valuable insight into the protein content under different stress and growth conditions that is indispensable for systems-level modeling.

Taken together, our results indicate that our fast and robust DIA-based approach can reliably and quantitatively retrieve the known differences and even reveal new and biologically meaningful regulation of protein expression, thereby providing a near comprehensive resource for yeast researchers and a valuable platform to support future studies in quantitative biology.

### Global regulation of the yeast proteome during glucose starvation and recovery

To gain better insights into how yeast regulate metabolism in response to a change in carbon source, we next expanded our analysis to investigate glucose starvation and glucose recovery. Yeast cultures were first grown to logarithmic phase in glucose, then switched to media containing ethanol as a non-fermentable carbon source. Following 19 hours growth in ethanol, glucose was replenished and the yeast were allowed to recover for 30 minutes or 2 hours (**Figure 3A**). Our workflow quantified 3,602 distinct proteins in total (**Supplementary Table 3**). The first PCA component segregated the growth conditions, with glucose being largely separated from the ethanol and recovery conditions (**Figure 3B, S3A**). To further investigate the regulation of metabolism in alternate carbon sources, we compared the proteome changes with those of the transcriptome. PCA analysis of the transcriptome also showed that the first component separated the growth conditions. Interestingly, in this case cells grown in ethanol were largely separated from the glucose (never starved) and glucose recovery conditions (**Figure 3C, S3B**), suggesting that during this metabolic shift, yeast cells remodel their gene expression first through rapid changes in transcription, which facilitates production of new proteins, and then remove proteins that are no longer required.

**Figure 3.**
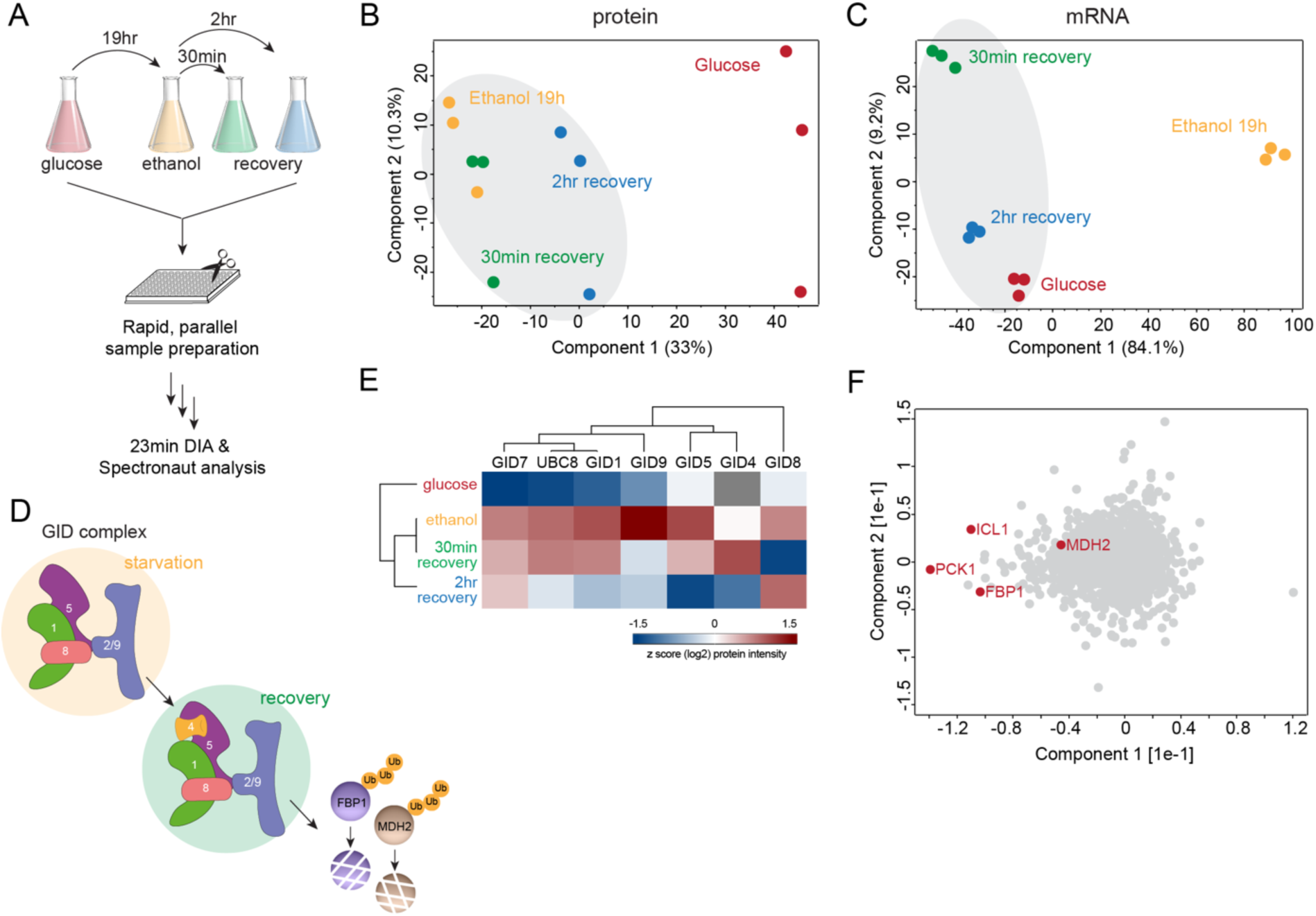
Global proteome changes of yeast under glucose starvation and recovery. **A**. Rapid yeast proteome profiling under glucose starvation and recovery. **B-C**. PCA plot of growth conditions along with their biological replicates based on their protein expression (B) and mRNA abundance (C) profiles. **D**. The GID E3 ubiquitin ligase is a key regulator of the switch from gluconeogenic to glycolytic growth as it degrades the gluconeogenic enzymes, including Fbp1 and Mdh2. **E**. Heat map of z-scored protein abundances (log2) of the GID complex subunits under glucose starvation and recovery in wildtype yeast cells. **F**. PCA plot of proteins during glucose starvation and recovery. Proteins marked in red represent the known GID complex substrates.

Several regulatory mechanisms contribute to carbohydrate metabolism, including allosteric regulation, reversible enzyme inactivation through covalent modifications, and irreversible loss of enzyme activity through proteolysis (reviewed in (57)). Importantly, we observed that protein turnover during glucose recovery occurs rapidly and in less than one cell division, suggesting an active mechanism of protein degradation. One such mechanism that has been well-characterized by our group and others is the ubiquitination and degradation of gluconeogenic enzymes by the GID (*<u>g</u>*lucose-*<u>i</u>*nduced *<u>d</u>*egradation) E3 ubiquitin ligase. The GID ligase is present at low levels in all growth conditions. However, during growth in ethanol, most of the GID subunits are induced leading to the formation of a yet inactive anticipatory complex: GID^Ant^. Following glucose replenishment, the substrate receptor, Gid4, is rapidly induced and joins the complex (40), allowing the recognition and subsequent degradation of the gluconeogenic proteins Fbp1, Mdh2, Icl1, and Pck1 via the Pro/N-degron pathway (**Figure 3D**) (38, 58-60). Indeed, our analysis confirmed that most components of the GID E3 ligase are upregulated around 4-fold during growth in ethanol, with the exception of Gid4, which is rapidly and transiently upregulated within 30 minutes of glucose replenishment (**Figure 3E**).

Intriguingly, PCA analysis of individual proteins revealed that the known substrates of the GID E3 ubiquitin ligase, Fbp1, Pck1, Icl1 and, to a lesser extent, Mdh2 are the major contributors to the segregation based on growth condition (**Figure 3F**). While the GID E3 ligase is known to be an important contributor to the regulation of yeast metabolism during the switch from gluconeogenic to glycolytic conditions, and is thought to have additional substrates, the lack of an obvious phenotype in GID mutants has made the identification of further substrates challenging. Thus, we applied our rapid and robust DIA-based methodology to search for novel regulatory targets of the GID E3 ligase.

### Identifying novel GID ligase-dependent regulation during recovery from carbon starvation

The structure and molecular mechanism of the GID E3 ligase are known, but the pathways it regulates are only beginning to be elucidated (38, 40, 61). While the role of the GID ligase in the regulation of gluconeogenesis is well-characterized, the conservation of this multi-protein complex throughout eukaryotes suggests that it likely regulates additional pathways. For example, the GID/CTLH complex has a role in erythropoiesis and spermatogenesis in human cells, and in embryogenesis in Drosophila (62-65). Thus, we set out to uncover additional pathways regulated by the GID E3 ligase by utilizing a combination of mutants. First, we used a deletion of the substrate receptor, Gid4, which targets proteins with either an N-terminal proline or a proline at position 2 via the Pro/N-degron pathway (38, 59, 60, 66). Deletion of Gid4 therefore should prevent substrate binding to the GID complex, and thereby inhibit degradation. However, Gid4, while conserved in human cells, is not conserved throughout all eukaryotes. For example, the GID complex in Drosophila lacks an identifiable Gid4 homolog (65), suggesting an alternate mode of recognition. In addition, in yeast, the protein Gid10 is an alternate substrate receptor of the GID complex (67), although no Gid10-dependent cellular substrates have been identified to date. To identify pathways regulated by the GID complex by a novel recognition pathway, we used a structure-based point mutant in the RING-domain-containing subunit, Gid2^K365A^, which eliminates catalytic activity without altering folding or complex assembly (40).

We compared the transcriptomes and proteomes of wildtype yeast to yeast containing either a Gid4 deletion or a Gid2 mutant (Gid2^K365A^) grown under the glucose starvation and recovery conditions described previously. Each condition was measured in triplicates using the rapid DIA method (**Figure 4A, Supplementary Table 3**). Importantly, there were no GID-dependent differences in mRNA levels following glucose replenishment (**Figure S4A**), demonstrating that the GID E3 ligase does not regulate protein synthesis, but rather the fate of existing proteins. To confirm that our DIA-based approach would be able to recognize bona fide GID substrates, we first examined the expression patterns of the well-characterized substrates Fbp1 and Mdh2. Indeed, in wildtype cells, Fbp1 and Mdh2 proteins are induced during growth in ethanol and then turned over within two hours of glucose recovery, with Fbp1 and Mdh2 protein levels reduced by around 8-fold and 5.7-fold, respectively. As expected, both proteins are also stabilized in the GID4-deleted and gid2-mutant cells (**Figure 4B, C**), confirming that we can robustly identify changes in expression of known substrates.

**Figure 4:**
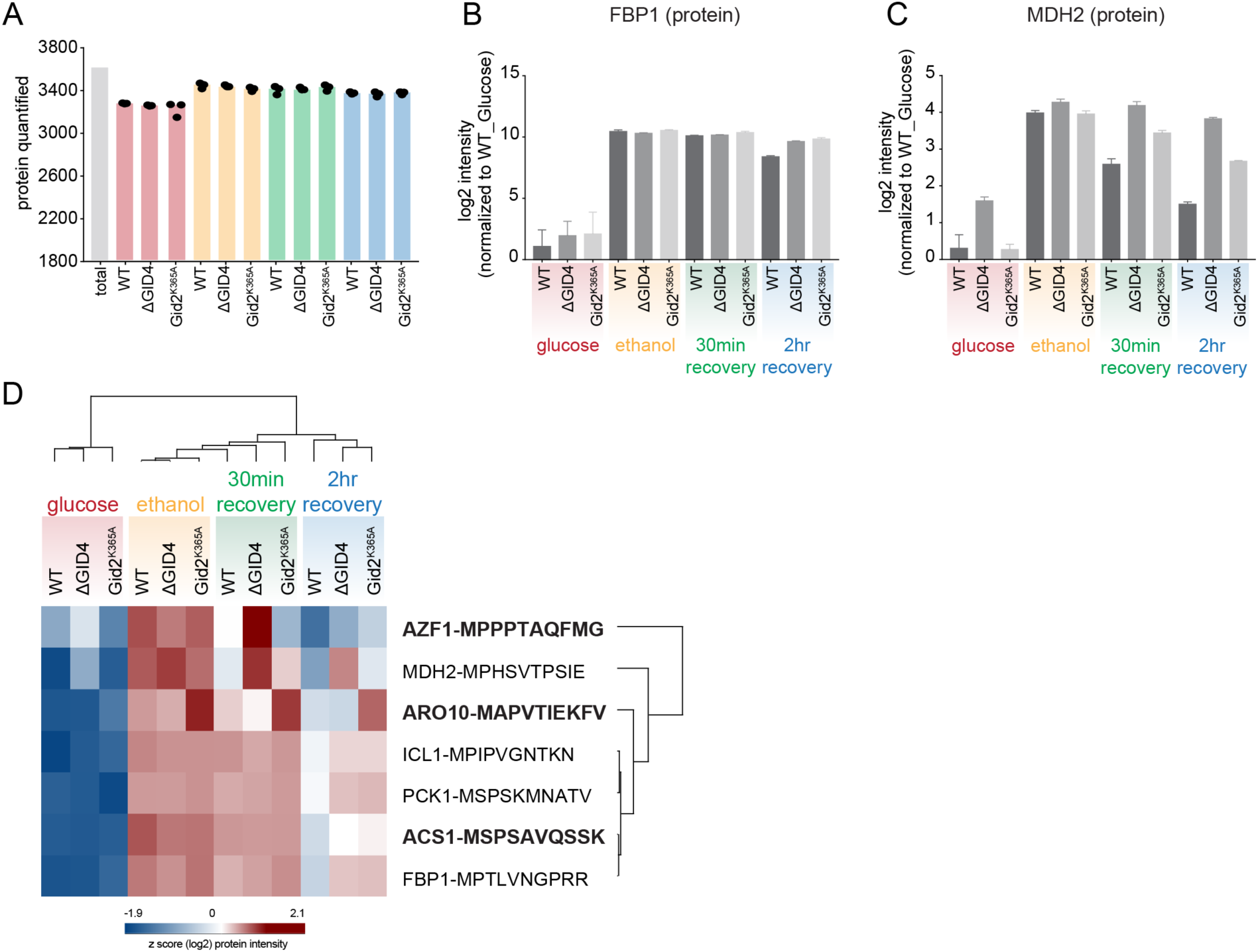
Our rapid and robust DIA-based approach identifies novel GID substrates under recovery after glucose starvation. **A**. Number of quantified proteins in WT, ΔGID4 and Gid2^K365A^ during glucose starvation and recovery. **B-C**. Bar graph shows abundances (log2) of Fbp1 (B) and Mdh2 (C) proteins that are normalized to wildtype Glucose (never starved) condition in WT, ΔGID4 and Gid2^K365A^ during glucose starvation and recovery. **D**. Heat map of z-scored protein abundances (log2) of the proteins which have the criteria of GID substrates.

To identify novel targets, we searched for proteins with an expression profile similar to the known substrates based on the following criteria: 1) the protein should be expressed more highly in ethanol than glucose, 2) its levels should decrease during glucose replenishment, and 3) after two hours of glucose replenishment, it should have a higher expression level in the GID4-deleted and/or gid2-mutant cells, compared to wildtype (**Figure S4B**). This provided a list of 31 proteins, including all four known GID substrates (Fbp1, Mdh2, Pck1, and Icl1) (**Figure S4C**). To further prioritize candidates, we limited our search to proteins with an N-terminal proline or a proline in the second position, a genetic and structural requirement of all known cellular substrates (38, 40, 60). The resulting list of 7 proteins consisted of the four known substrates, the transcription factor Azf1, and the metabolic enzymes Aro10 and Acs1 (**Figure 4D**). Interestingly, Azf1 has already been implicated in regulation of GID4 transcription (68), suggesting its upregulation in the GID deficient cells may be a cellular compensation mechanism. However, because we did not observe any GID-dependent mRNA expression changes (**Figure S4A**), we eliminated Azf1 from further analysis. Acs1 was significantly stabilized in both the GID4-deleted and gid2-mutant cells, whereas Aro10 was only significantly stabilized in the gid2-mutant.

In order to validate Aro10 and Acs1 as novel GID targets *in vivo*, we used the promoter reference technique (38, 69), a transcription independent method to examine protein turnover. In this method, yeast cells are transformed with a plasmid expressing the test substrate and the control protein DHFR from identical promoters (**Figure 5A**). The transcribed products carry tetracycline-binding RNA aptamers which inhibit protein expression at the level of translation following the addition of tetracycline to the media, allowing the fate of the existing protein to be monitored. Importantly, this method selectively terminates synthesis of our test proteins, thus the induction of Gid4 and activation of the GID complex is not impaired. In agreement with our proteomic findings, the Acs1 protein is completely stabilized in both GID2-and GID4-deleted cells (**Figure 5B**), while the Aro10 protein is stabilized in GID2-deleted but not GID4-deleted cells (**Figure 5C**), indicating a potential novel, Gid4-independent regulation. Thus, Acs1 and Aro10 are confirmed to be regulatory targets of the GID ligase during the switch from gluconeogenic to glycolytic conditions.

**Figure 5.**
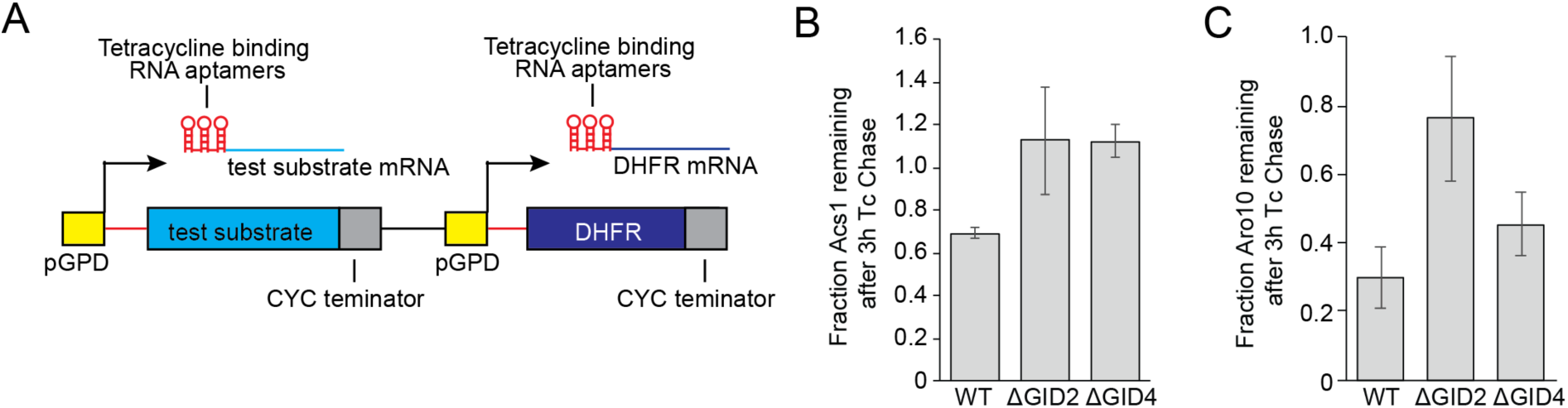
*In vivo* validation of novel GID targets. **A**. Schematic of promoter reference technique. **B-C**. Quantification of Acs1 (B) and Aro10 (C) degradation, based on at least 4 independent replicates. Bars represent mean values, and error bars represent standard deviation

## DISCUSSION

Here, we described a straightforward, streamlined and reproducible systems biology approach for yeast proteome profiling using data-independent acquisition (DIA) to analyze biological pathways much faster and with greater depth. Our minimalistic workflow employs plate-based preparation of digested yeast cell lysate, requires only a few micrograms of yeast as input, and no labeling or special equipment, making it especially amenable for application in non-specialized research groups. Despite its simplicity, it robustly and quantitatively profiles hundreds of largely covered yeast proteomes (80% of the expressed proteome at normal growth conditions (7) within an unprecedented throughput (100 samples in ∼2.2 days).

The ability of cells to adapt to stress or changes in environmental conditions relies on extensive proteome remodeling (70-75). Understanding these changes provides broad insight into the molecular mechanisms underlying many processes including, heat stress, adaptation to nutrient availability, and regulation of cell division. Applying our DIA-based approach to profile protein levels in systems-wide response to several stress and growth conditions demonstrated its robustness and specificity. In addition, our work provides an in-depth resource on stress mediators regulated at the protein level, which will complement the widely available yeast transcriptome data, and further allow yeast researchers to probe numerous biological pathways of interest, including stress response pathways, autophagy, and nutrient signaling pathways.

In addition to identifying proteome changes during stress, we applied our DIA-based systems biology approach to discover novel proteins that are regulated by the GID E3 ubiquitin ligase, a key regulator in the switch from gluconeogenic to glycolytic conditions (54, 61, 76). Despite the importance of the GID complex in metabolic regulation, identification of additional substrates has been hindered by the lack of an obvious phenotype, variable kinetics of protein degradation, and the necessity for a sensitive read-out. Our generic and unbiased approach, however, robustly identified two novel proteins regulatory targets of the GID complex: Acs1 and Aro10, further highlighting the importance and need for quantitative proteome datasets to provide a basis for functional studies.

Interestingly, both Acs1 and Aro10, while not considered gluconeogenic enzymes, are important regulators of metabolism and cellular respiration during anaerobic growth. Acs1 encodes one of two isoforms of yeast Acetyl-CoA synthetase, which catalyzes the formation of acetyl-CoA from acetate and CoA. Acs1 has a much higher affinity for acetate than its isoform Acs2, making it more desirable for acetyl-CoA production when acetate is limiting, as it is the case during growth on non-fermentable carbon sources (77). During glycolytic growth, however, the main energy flux does not require Acs1/2 function, Acs1 expression is suppressed and existing Acs1 protein must be degraded. Aro10 encodes a phenylpyruvate decarboxylase that catalyzes an irreversible step in the Ehrlich pathway, which provides a more energetically favorable means of NADH regeneration during anaerobic growth. Following glucose replenishment, NADH is regenerated through glycolysis, and thus Aro10 function is no longer required (78, 79).

Here, we show that both Acs1 and Aro10 turnover are dependent on the catalytic activity of the GID complex, via its RING-containing subunit, Gid2. Intriguingly, only Acs1 turnover is dependent on the well-characterized substrate receptor, Gid4, suggesting an alternate mode of recognition for Aro10. Indeed, an additional substrate receptor, Gid10, has recently been identified (40, 67), raising the possibility that Aro10 may be the first substrate identified in this novel recognition pathway. Alternatively, Aro10 recognition may be facilitated by a yet-to-be identified substrate receptor, or an alternative mechanism. In either case, the regulation of Aro10 suggests that the GID ligase may function with separable catalytic and substrate recognition elements, a mechanism, previously described for SCF (Skp1-Cullin-Fbox) E3 ligases (80, 81) that provides a flexible means for linking a single E3 to a greater number of substrates. Intriguingly, expression of the GID substrate receptors is induced during several other cellular stresses, including osmotic shock, heat shock, and nitrogen starvation (40, 67, 71), suggesting that the GID complex may play an important role in rewiring metabolic pathways during adaptation to a wide variety of stress conditions.

Taken together, the GID-dependent regulation of Acs1 and Aro10, along with the previously known substrates, suggest that the GID complex is a multifunction metabolic regulator that influences multiple cellular pathways simultaneously to allow for an efficient switch from gluconeogenic to glycolytic conditions. Moreover, our findings demonstrate that our DIA-based systems biology approach is capable of simultaneously identifying changes to multiple cellular pathways which are integrated to maintain cellular homeostasis. While we here identified specific targets of an E3 ligase, our workflow can be readily adopted by the community to probe numerous cellular pathways, including kinase signaling pathways or cell-cycle dependent changes. Furthermore, its speed allows the analyzing of at least 15 conditions, in triplicate, per day, making it particularly well suited for screens. For example, the effect of each of the ∼80 yeast E3 ligases on the global proteome could be ascertained in just five days, or each of the ∼117 yeast kinases in about one week. In addition, our workflow can be easily adapted to identify changes in post-translational modifications when coupled with an enrichment step.

Thus, the speed and reproducibility of the DIA-based approach presented here allows researchers to probe complex biological pathways and identify novel regulatory mechanisms. We are currently integrating the Evosep HPLC system into our approach as it eliminates the overhead time between sample pick up and start of MS measurement by using pre-formed gradients (82). Radically simplified workflows like the one described here could be extended to other organisms, generating high-quality quantitative proteome datasets which are required to explain biological processes on a system-wide level (83, 84). Given that the expressed human proteome (around 15,479 proteins, https://www.proteomicsdb.org) is only around three times larger than the expressed yeast proteome (5,391 proteins, (45)), with only three fold increase in proteomic depth, we anticipate fast single run DIA approaches will also be suitable for analysis of the human proteome.

## MATERIALS AND METHODS

### Yeast Strains and Growth Conditions

All yeast strains used in this study are derivatives of BY4741 and are listed in Table 1. For rich conditions, yeast cultures were grown in YPD (1% yeast extract, 2% peptone, 2% glucose) or SD (0.67% yeast nitrogen base without amino acids, 2% glucose, containing the appropriate amino acid supplements) media. Unless otherwise specified, yeast were grown at 30°C. For heat shock conditions, yeast cultures were grown in YPD to an OD_600_ of 1.0 and then shifted to the indicated temperature for 1 hour. For osmotic shock conditions, yeast cells were grown in YPD to and OD_600_ of 1.0, pelleted at 3,000rpm for 3 minutes, and resuspended at an OD_600_ of 1.0 in pre-warmed YPD + 0.5M NaCl. For glucose starvation, yeast cells were grown in YPD to an OD_600_ of 1.0-2.0, pelleted at 3,000rpm for 3 minutes, washed once with YPE (1% yeast extract, 2% peptone, 2% ethanol), resuspended in pre-warmed YPE at an OD_600_ of 1.0, and grown at 30°C for 19 hours. For glucose recovery, yeast cells were pelleted after 19 hours growth in YPE, and resuspended to an OD_600_ of 1.0 in YPD, and allowed to grow at 30°C for 30 minutes or 2 hours. For amino acid starvation, yeast cells were grown in SC to an OD_600_ of 1.0-2.0, pelleted at 3,000rpm for 3 minutes, washed once with SD-AA (0.67% yeast nitrogen base without amino acids, 2% glucose, 20mg/L uracil), resuspended in SD-AA to an OD_600_ of 1.0, and allowed to grow for one hour. For nitrogen depletion, yeast cells were grown in SC to an OD_600_ of 1.0-2.0, pelleted at 3,000rpm for 3 minutes, washed once with SD-N (0.17% yeast nitrogen base without amino acids or ammonium sulfate, 2% glucose), resuspended in SD-N to an OD_600_ of 1.0, and allowed to grow for one hour. For proteomics analysis, 50 ODs of cells were pelleted at 3,000rpm for 3 minutes, flash frozen in liquid nitrogen, and stored at −80°C until lysis. For transcriptomes analysis, 10 ODs of yeast were pelleted, flash frozen in liquid nitrogen, and stored at −80°C.

### Protein Degradation Assays (promoter reference technique)

Protein degradation assays using the promoter reference technique were done as previously described (69). Cells were transformed with plasmid expressing a test substrate and DHFR fromidentical promoters containing tetracycline-repressible RNA-binding elements. Yeast cells were then grown in SC media lacking histidine, starved in SE (2% ethanol) media lacking histidine for 19 hours, and then allowed to recover for the indicated times in SC media lacking histidine. At each time point, 1.0 ODs of yeast cells were pelleted, flash frozen in liquid nitrogen, and stored at −80°C until lysis.

For lysis, yeast cells were resuspended in 0.8mL of 0.2M NaOH, followed by incubation on ice for 20 minutes, and then pelleted at 11,200xg for 1 minute. The supernatant was removed and the pellet resuspended in 50uL HU buffer, and incubated at 70°C for 10 minutes. The lysate was precleared by centrifugation at 11,200xg for 5 minutes, and then loaded onto a 12% SDS-PAGE gel. Protein samples were transferred to nitrocellulose membrane, and then visualized by western blot using αFLAG (Sigma, F1804) and αHA (Sigma H6908) primary antibodies, and Dylight 633 goat anti-Mouse (Invitrogen 35512) and Dylight 488 goat anti-rabbit (Invitrogen 35552) secondary antibodies. Membranes were imaged on a typhoon scanner (Amersham). Bands were quantified with ImageStudio software (Licor).

### mRNA sequencing

Harvested and frozen cells were sent to Novogene Co., Ltd (Hong Kong) for RNA extraction, library preparation, mapping, and bioinformatics analysis. Briefly, 3μg of RNA was used for library generation uding NEB Next Ultra RNA Library Prep Kit for Illumina (NEB). The library preparations were sequences on an Illumina Hi-Seq platform and 125bp/150bp paired-end reads were generated. Reads were indexed using Bowtie v2.2.3 and paired-end clean reads were aligned to the reference genome using TopHat v2.0.12. HTSeq v0.6.1 was used to count the read numbers mapped to each gene, and then FPKM (expected number of Fragments Per Kilobase of transcript sequence per Millions base pairs sequenced) of each gene was calculated based on the length of the gene and read counts mapped to the gene. The transcriptome data analysis was performed as explained in “Data processing and bioinformatics analysis”.

### Sample preparation for MS analysis

SDC lysis buffer (1 % SDC and 100 mM Tris pH 8.4) were added to the frozen cell pellets to achieve a protein concentration of ∼2–3 mg per ml. Lysates were immediately heat-treated for 5 minutes at 95 °C to facilitate lysis and to inactivate endogenous proteases and transfreed to a 96-well plate. Lysates were next homogenized with sonication. Protein concentrations were estimated by tryptophan assay (27) and then all samples were diluted to equal protein concentrations in a 96-well plate. To reduce and alkylate proteins, samples were incubated for 5 minutes at 45°C with CAA and TCEP, final concentrations of 10 mM and 40 mM, respectively. Samples were digested overnight at 37°C using trypsin (1:100 w/w, Sigma-Aldrich) and LysC (1/100 w/w, Wako). The following day, peptide material was desalted using SDB-RPS StageTips (Empore) (27). Briefly, samples were diluted with 1% TFA in isopropanol to a final volume of 200 µl and loaded onto StageTips, subsequently washed with 200 µl of 1% TFA in isopropanol and 200 µl 0.2% TFA/ 2% ACN. Peptides were eluted with 80 µl of 1.25% Ammonium hydroxide (NH4OH)/ 80% ACN and dried using a SpeedVac centrifuge (Eppendorf, Concentrator plus). Samples were resuspended in buffer A* (0.2% TFA/ 2% ACN) prior to LC-MS/MS analysis. Peptide concentrations were measured optically at 280nm (Nanodrop 2000, Thermo Scientific) and subsequently equalized using buffer A*. 300ng peptide was subjected to LC-MS/MS analysis.

To generate the spectral library for DIA measurements cells were lysed in SDC buffer, followed by sonication, protein quantification, reduction, and alkylation and desalting using SDB-RPS cartridges (see above). Around 8 or 30 µg of peptides were fractionated into 8 or 24 fractions, respectively, by high pH reversed-phase chromatography as described earlier (85). Fractions were concatenated automatically by shifting the collection tube during the gradient and subsequently dried in a vacuum centrifuge and resuspended in buffer A*.

### LC-MS/MS Measurements

Samples were loaded onto a 20 cm reversed phase column (75 μm inner diameter, packed in house with ReproSil-Pur C18-AQ 1.9 μm resin [Dr. Maisch GmbH]). The column temperature was maintained at 60°C using a homemade column oven. A binary buffer system, consisting of buffer A (0.1% formic acid (FA)) and buffer B (80% ACN plus 0.1% FA), was used for peptide separation, at a flow rate of 450 nl/min. An EASY-nLC 1200 system (Thermo Fisher Scientific), directly coupled online with the mass spectrometer (Q Exactive HF-X, Thermo Fisher Scientific) via a nano-electrospray source, was employed for nano-flow liquid chromatography. We used a gradient starting at 5% buffer B, increased to 35% in 18.5 minutes, 95% in a minute and stayed at 95% for 3.5 minutes. The mass spectrometer was operated in Top10 data-dependent mode (DDA) with a full scan range of 300-1650 m/z at 60,000 resolution with an automatic gain control (AGC) target of 3e6 and a maximum fill time of 20ms. Precursor ions were isolated with a width of 1.4 m/z and fragmented by higher-energy collisional dissociation (HCD) (NCE 27%). Fragment scans were performed at a resolution of 15,000, an AGC of 1e5 and a maximum injection time of 60 ms. Dynamic exclusion was enabled and set to 30 s. For DIA measurements full MS resolution was set to 120,000 with a full scan range of 300-1650 m/z, a maximum fill time of 60 ms and an automatic gain control (AGC) target of 3e6. One full scan was followed by 12 windows with a resolution of 30,000 in profile mode. Precursor ions were fragmented by stepped higher-energy collisional dissociation (HCD) (NCE 25.5, 27,30%).

### Data processing and bioinformatics analysis

Spectronaut version 13 (Biognosys) was used to generate the spectral libraries from DDA runs by combining files of respective fractionations using the yeast FASTA file (Swissprot, 2018). For the generation of the proteome library default settings were left unchanged. DIA files were analyzed using the proteome library with default settings and enabled cross run normalization. The Perseus software package versions 1.6.0.7 and 1.6.0.9 and GraphPad Prism version 7.03 were used for the data analysis (86). Protein intensities and mRNA abundances were log2-transformed for further analysis. The data sets were filtered to make sure that identified proteins and mRNAs showed expression or intensity in all biological triplicates of at least one condition and the missing values were subsequently replaced by random numbers that were drawn from a normal distribution (width=0.3 and down shift=1.8). PCA analysis of stress and growth conditions and biological replicates was performed as previously described in (87). Multi-sample test (ANOVA) for determining if any of the means of stress and growth conditions were significantly different from each other was applied to both mRNA and protein data sets. For truncation, we used permutation-based FDR which was set to 0.05 in conjunction with an S0-parameter of 0.1. For hierarchical clustering of significant genes and proteins, median protein or transcript abundances of biological replicates were z-scored and clustered using Euclidean as a distance measure for row clustering. GO annotations were matched to the proteome data based on Uniprot identifiers. Annotation term enrichment was performed with either Fisher exact test or 1D tool in Perseus. Annotation terms were filtered for terms with an FDR of 5% after Benjamini-Hochberg correction.

## Data availability

All data supporting findings of this study are available within this article and in supplementary tables.

## ACKNOWLEDGEMENTS

This work was supported by the Max-Planck Society for the Advancement of Science. We thank Arno Alpi, Florian Meier, Igor Paron, Christian Deiml, Johannes B. Mueller, Viola Beier, Kirby Swatek and all the members of the departments of Proteomics and Signal Transduction and Molecular Machines and Signaling at Max-Planck-Institute of Biochemistry for their assistances and helpful discussions.

## SUPPLEMENTARY FIGURES

**Supplementary Figure 1.**
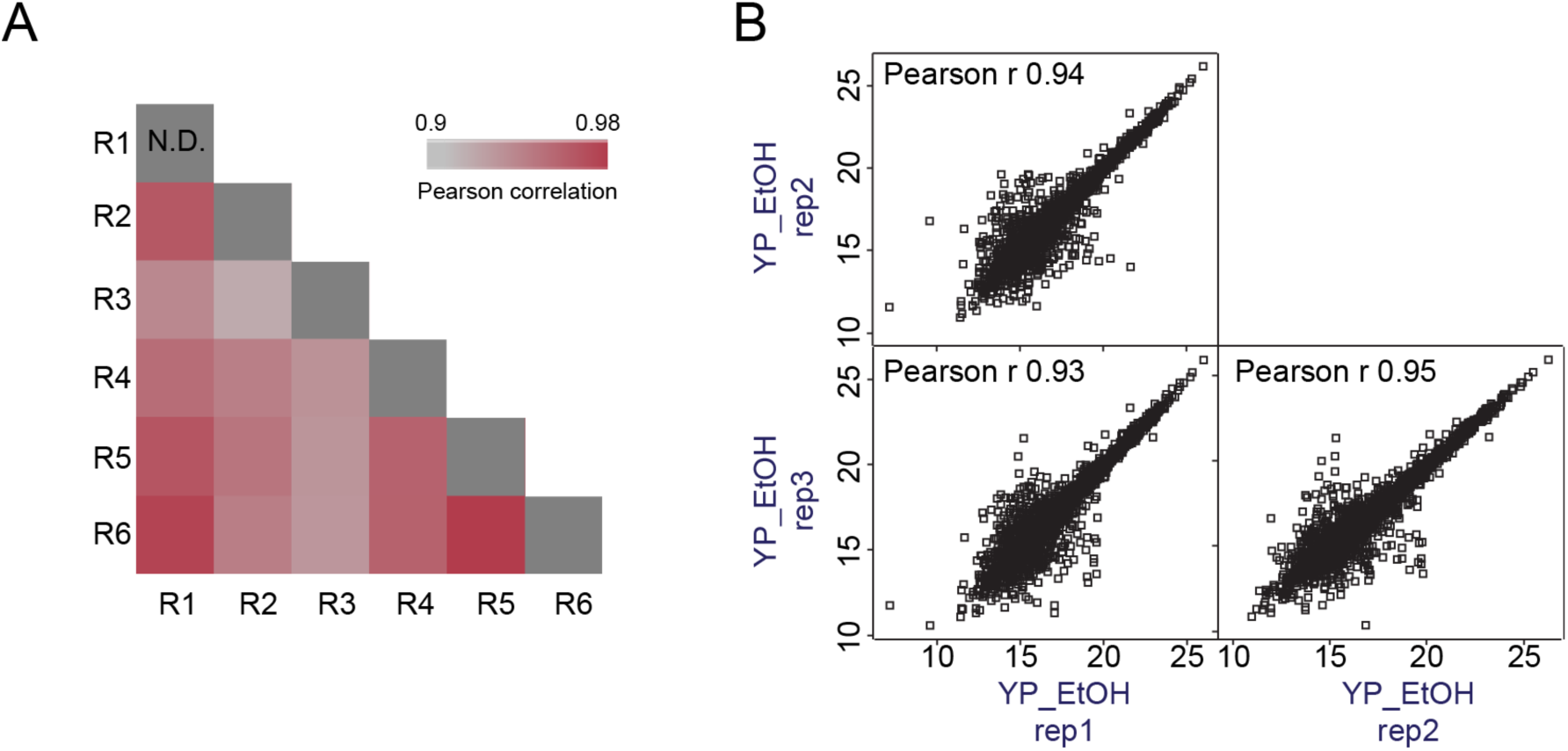
Reproducibility of the DIA-based workflow. A. Correlation based clustering illustrating the reproducibility between workflow replicates. High (0.98) and lower (0.9) Pearson correlations are denoted in red and grey, respectively. B. The correlation plots illustrating the reproducibility between biological replicates in the yeast stress experiment. Pearson correlation coefficients are shown in the upper left corner.

**Supplementary Figure 2.**
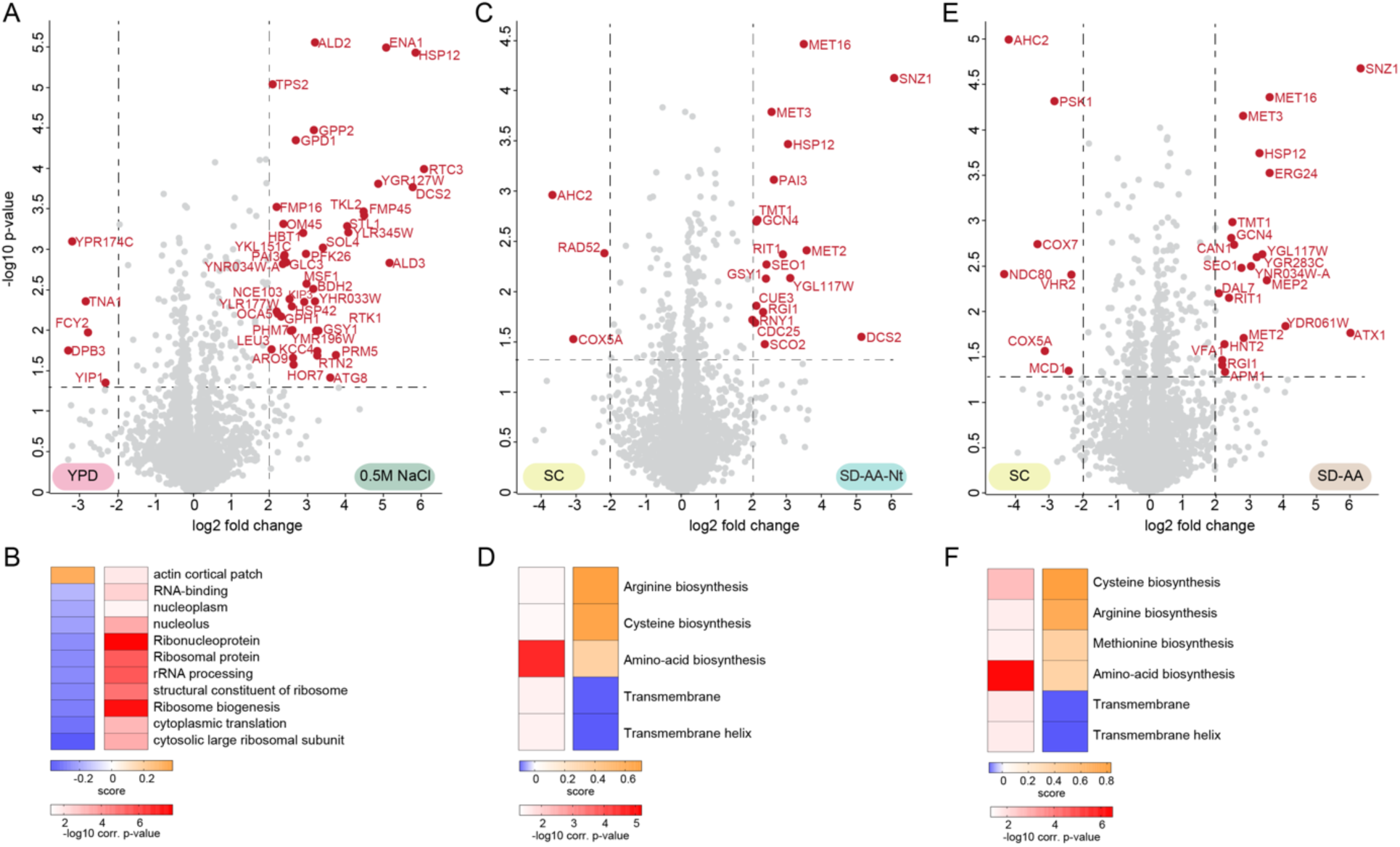
Mapping changes to the yeast proteome in response to osmotic shock, and amino acid and nitrogen depletion. Volcano plot of the (-log10) p-values vs. the log2 protein abundance differences between 0.5M NaCl (osmotic shock) vs. YPD (A), SD-AA-Nt vs. SC (C), and SD-AA vs. SC (E). The significant proteins (red dots) are determined based on p-value < 0.05 and at least 4-fold change on both sides. GO-term enrichment in the 0.5M NaCl (osmotic shock) vs. YPD (B), SD-AA-Nt vs. SC (D), and SD-AA vs. SC (F). fold change dimension (1D enrichment, FDR < 5%). Terms with positive enrichment scores are enriched in stress condition over YPD or SC control and vice versa.

**Supplementary Figure 3.**
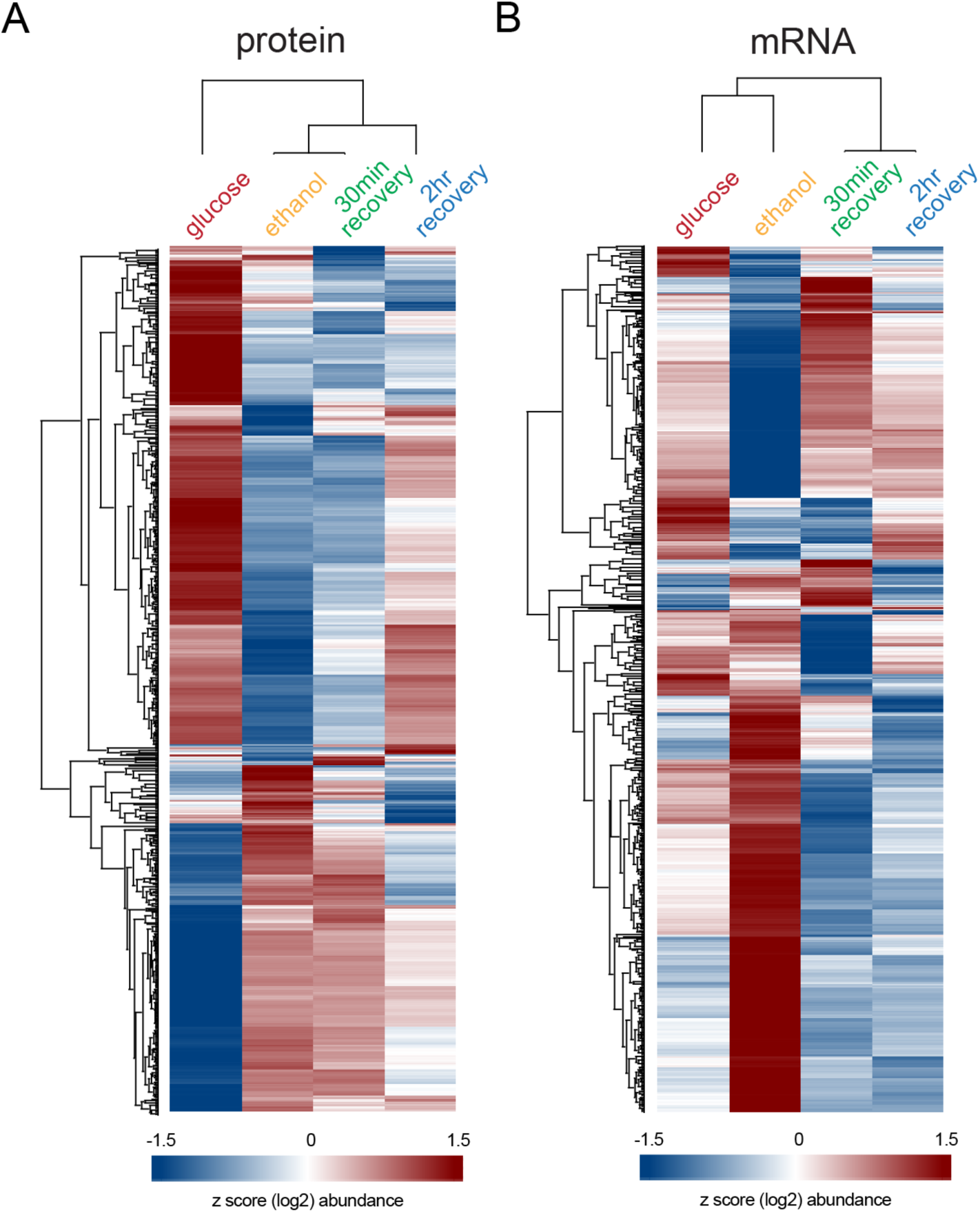
Global proteome and transcriptome changes of yeast under glucose starvation and recovery. A-B. Heat map of z-scored and differentially regulated proteins (A) and mRNAs (B) (log2) in wildtype yeast during glucose starvation and recovery.

**Supplementary Figure 4.**
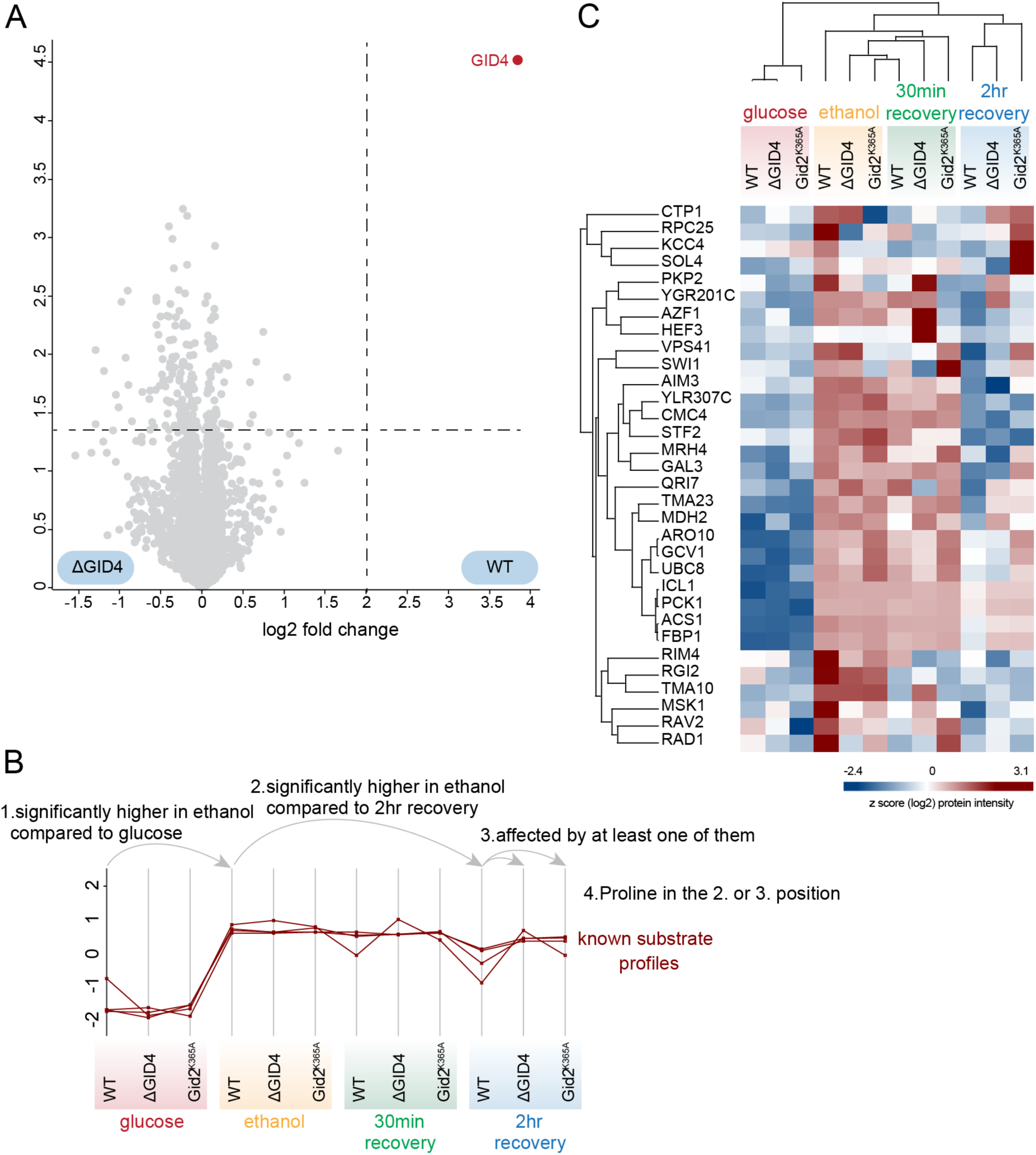
Identifying GID ligase targets during recovery from carbon starvation. A. Volcano plot of the (-log10) p-values vs. the log2 mRNA abundance differences between wildtype vs. Gid4 (substrate receptor) deletion. GID4 (shown in red) was the only significant hit based on p-value < 0.05 and at least 4-fold change on both sides. B. The criteria of the known GID substrates based on their protein profiles: (1) the protein is expressed significantly higher in ethanol compared to glucose and (2) 2hr recovery, (3) decreased abundance of the protein during recovery is dependent on the GID complex, and (4) having Proline (N-degron) in position 2 or 3. C. Heat map of z-scored potential GID targets which meet the first three conditions in panel B during glucose starvation and recovery.

